# Deep learning enables fast, gentle STED microscopy

**DOI:** 10.1101/2023.01.26.525571

**Authors:** Vahid Ebrahimi, Till Stephan, Jiah Kim, Pablo Carravilla, Christian Eggeling, Stefan Jakobs, Kyu Young Han

## Abstract

STED microscopy is widely used to image subcellular structures with super-resolution. Here, we report that denoising STED images with deep learning can mitigate photobleaching and photodamage by reducing the pixel dwell time by one or two orders of magnitude. Our method allows for efficient and robust restoration of noisy 2D and 3D STED images with multiple targets and facilitates long-term imaging of mitochondrial dynamics.

## Main text

Stimulated emission depletion microscopy (STED)^1,2^ is a super-resolution fluorescence imaging technique that can reveal biological structures in live cells with greater than 50 nm resolution^3^. Hereby, the effective fluorescence area is confined to nanoscales by overlapping the diffraction-limited excitation spot with a fluorescence-depleting spot exhibiting a central intensity of zero (such as a doughnut-shaped spot). Increasing the intensity of the depletion beam leads to an increase in resolution but often causes adverse effects such as photobleaching^4^ and phototoxicity^5^, preventing long-term monitoring of samples. Although optimized optical parameters^2^, multiple off states^6^, exchangeable fluorophores^7,8^, or sophisticated illumination^9^ and data acquisition schemes^10,11^ can circumvent these problems to some extent, the improvement is often small, or the choice of fluorophores is limited. In principle, reducing the STED exposure time can decrease photodamage^5^; however, a short pixel dwell time results in a poor signal-to-noise ratio (SNR) and consequently degrades image resolution^12^.

Emerging deep learning approaches have proposed different solutions to address the tradeoffs between spatial/temporal resolution, SNR, and phototoxicity^13-15^. In particular, converting confocal images to high-resolution STED images called cross-modality image restoration has shown promising results^16,17^. Here, we show that denoising STED images for image restoration is advantageous over other deep learning methods and can significantly enhance the performance of STED microscopy, i.e., the increased imaging speed and the extended observation time.

We used a two-step prediction architecture based on a U-Net^18^, and a residual channel attention network (RCAN)^19^ (UNet-RCAN), in which a single U-Net restores the broad contextual information and an RCAN reconstructs the final super-resolution images (Fig.1a,Extended Data Fig.1). For training and testing, we acquired multiple pairs of low SNR STED images at a short pixel time (*Δt*) and high SNR STED images at a long pixel time. If necessary, a drift correction was applied for the registration of each pair (Extended Data Fig.2). Alternatively, we obtained high SNR images and generated low SNR counterparts by adding noise (Methods), which corresponded well with the results of sequentially acquired training set (Extended Data Fig.3). We generally experienced that 20 image data sets were enough to train our denoising algorithm.

**Fig. 1.**
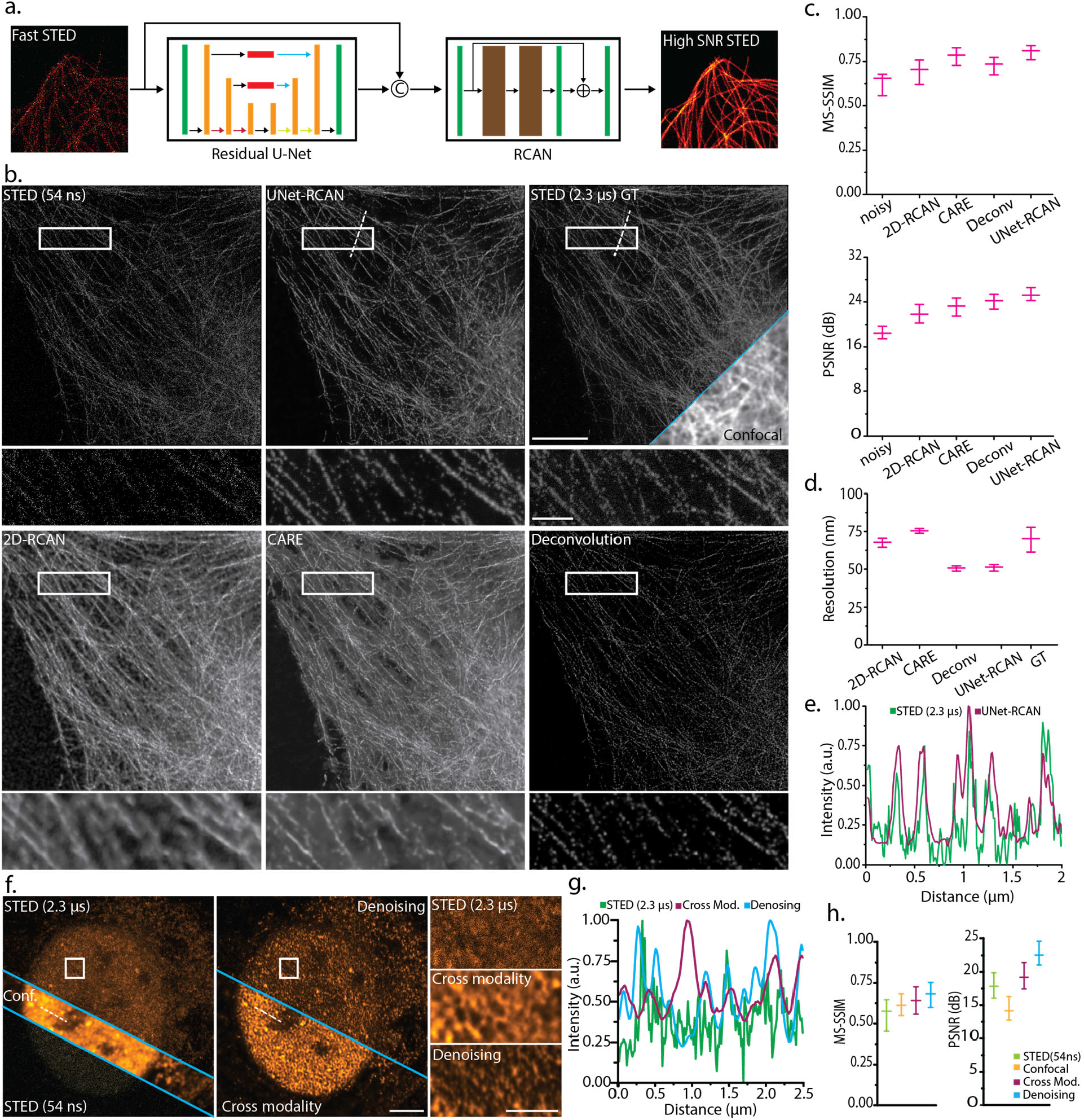
Restoration of noisy STED images by UNet-RCAN. (**a**) The network architecture of two-step denoising. (**b**) Denoising results of UNet-RCAN, 2D-RCAN, CARE, and deconvolution on noisy 2D-STED images (*Δt* = 54 ns) for β-tubulin (STAR635P) in U2OS cells in comparison to the ground-truth (GT) STED data (*Δt* = 2.3 μs). (**c**) Quantitative comparisons of the predicted results for the methods used in (b). Mean and standard deviation are displayed (*n* = 10). (**d**) Resolution analysis by phase decorrelation for the predicted results. Mean and standard error of mean are displayed (*n* = 10). (**e**) Line profiles along the white dashed lines in (b). (**f**) Comparison of cross-modality and denoising methods for restoring high SNR STED images. Cross-modality used confocal images as input. Histone markers (H3K9ac) were labeled with Atto647N in U2OS cells. (**g**) Line profiles along the white dashed lines in (f). (**h**) Quantitative metrics of the predicted results by cross-modality and denoising (STED = 54 ns). Scale bars, 5 μm (b,f) and 1 μm for their magnified regions.

To investigate the performance of our network on denoising STED microscopy data, we obtained 2D-STED images of microtubules (β-tubulin) labeled with STAR635P in fixed U2OS cells. The pixel times of noisy input and ground-truth were 0.054 μs and 2.3 μs, respectively. We can clearly see significant improvement in SNR by comparing the predicted images to the noisy STED data (Fig. 1b), indicating our approach can reduce the pixel time of STED microscopy by >40-fold. Compared to other networks or deconvolution, our approach yields improved accuracy of predictions in terms of multi-scale structural similarity index (MS-SSIM) and peak SNR (PSNR) (Fig.1c). Importantly, our method maintains the lateral resolution of STED images assessed by decorrelation analysis (Fig.1d). For example, for the ground-truth STED image we estimated a resolution of 76 ± 3 nm whereas the predicted result showed 50 ± 2 nm. It is likely that the resolution improvement by UNet-RCAN over ground-truth STED images is due to the increase of SNR. We also validated our approach on various subcellular targets (Extended Data Fig.4) and two-color samples (Extended Data Fig.5).

We found that our method is robust at different STED powers (Extended Data Fig. 6). The spatial resolution of the predicted results is consistent with the scaling law of STED microscopy. Although the SNR of STED images depends on numerous factors, including the excitation intensity, fluorophores, labeling density, pixel time, etc., we can estimate how reliable our prediction is given a certain level of SNR (Extended Data Fig.7). It is unavoidable that the image quality parameters somewhat drop for very low SNR images. We want to emphasize that our two-step deep learning approach has merit over others. Unlike content-aware image restoration (CARE)^13^, UNet-RCAN maintains the high spatial resolution of the STED images (Figs. 1d,e). Compared to cross-modality image restoration^16,17^ and deconvolution, our approach generates fewer artifacts in prediction, especially for low SNR images (Figs. 1f-h, Extended Data Fig.8 and Supplementary Note 1).

The low-exposure images used in our denoising method are significantly less susceptible to photobleaching. While the signal level halved after 5-10 frames in conventional STED imaging of β-tubulin (STAR635P) and histone (Alexa 594), our approach maintained the signal for over 300 frames (Figs. 2a,b). Our denoising approach also facilitates high-throughput STED imaging (Extended Data Fig.9). It took 21 min to record 744 STED images (2,048×2,048 pixels) over a 1.0× 0.78 mm^2^ region; it would take ∼14 hours to do comparable imaging using traditional STED.

**Fig. 2.**
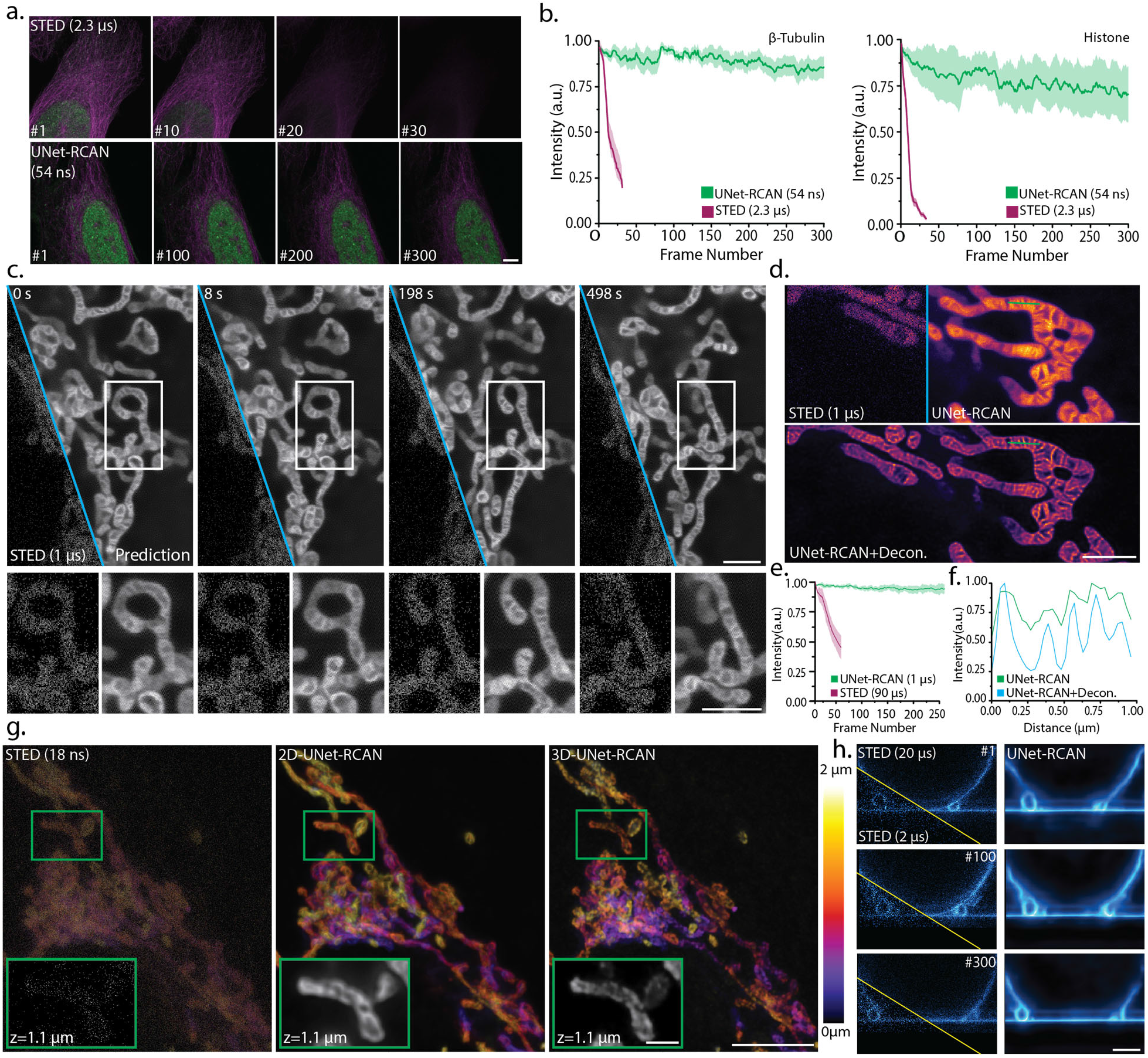
Reduced photobleaching and photodamage of STED imaging by UNet-RCAN. (**a**) Time-lapse STED images obtained by conventional STED (top, *Δt* = 2.3 μs) and UNet-RCAN using fast STED data (bottom, *Δt* = 0.054μs). β-tubulin (STAR635P, magenta) and histone (Alexa 594, green) were imaged in U2OS cells. (**b**) Photobleaching analysis of two-color STED images used in (a). Shaded areas are standard deviations of fluorescence intensities (*n* = 5). (**c**) Live-cell STED recording of mitochondrial cristae in HeLa cells (*Δt* = 1 μs) labeled with PK Mito Orange. (**d**) Denoising and deconvolved STED recording of mitochondrial cristae in COS-7 cells (*Δt* = 1 μs) labeled with PK Mito Orange (**e**) Fluorescence time traces of mitochondrial cristae in HeLa cells over 250 consecutive frames (2 s/frame). The fluorescence time trace of STED recording with *Δt* = 90 μs is displayed for comparison. (**f**) Line profiles along a dashed line in (d). (**g**) 2D- and 3D-UNet-RCAN prediction results for a noisy z-stack of 3D-STED imaging of TOM20 (Atto647N). The z pixel size is 65 nm. (**h**) Denoising results for *xz* time-lapse STED recording of membrane fusion dynamics (*Δt* = 2 μs) over 300 frames in comparison to ground-truth data (*Δt* = 20 μs). Scale bars, 5 μm (a,g), 2 μm (c,d,h) and 1 μm for the magnified regions in (g).

Next, we applied our denoising to live-cell STED imaging. Its gentle illumination (*Δt* = 1 μs) enabled us to capture >200 frames of STED images of mitochondrial dynamics in HeLa cells with minimal phototoxicity, which is a ten-fold increase compared to the conventional STED (*Δt* = 90 μs) (Figs.2c,e, Supplementary Videos 1-3). Our model preserved the shape of the original cristae images (Supplementary Figs.1,2), and deconvolution can further improve their resolution (Figs.2d,f, Supplementary Video 4).

It is straightforward to extend our approach to volumetric STED imaging. For this application, we trained a 3D UNet-RCAN using 3D stacks of STED images acquired by 2D or 3D STED (Fig.2g, Extended Data Fig.10). Our predicted results using fast 3D STED input images (*Δt* = 0.018 μs) clearly showed the hollow shape of mitochondria labeled to TOM20. The 3D model showed improvement in prediction accuracy compared to the 2D model, likely due to the effective consideration of noise^17^. The UNet-RCAN is also applicable to time-lapse 3D-STED *xz* imaging of fast dynamics. It realizes the disclosure of the fusion dynamics between a giant unilamellar vesicle and a supported bilayer^8^ with a temporal resolution of 315 ms/frame (*Δt* = 2 μs) compared to 3.15 s/frame (*Δt* = 20 μs) (Fig. 2h, Supplementary Video 5). It generally leads to noise-reduced data and even clearly recovering the membrane ghosts, a typical artifact for 3D STED images of membranes^20^.

In summary, denoising STED images is a powerful tool for fast, long-term super-resolution imaging. It is readily implementable without any hardware changes. When combined with other concepts like adaptive illumination^10^ and/or event-triggered imaging^21^, our method can further reduce the phototoxic effects of live-cell STED to a bare minimum. Similarly, our concept could be combined with an ultrafast scanning system^22^ to enable gentle live-cell nanoscopy at maximum speed.

## Methods

### Architecture of UNet-RCAN

We adopted a two-step prediction architecture from the multi-stage progressive image restoration (MPRNet)^23^, but it was modified for high-resolution fluorescence imaging as follows. The first subnetwork is a residual U-Net^18^, a convolutional neural network for image reconstruction through down-sampling and up-sampling operations like CARE^13^. An encoder consists of a residual convolution block — a first convolution layer, a leaky rectified linear unit (LeakyReLU; leakage factor = 0.3) as an activation function, and a second convolution layer, followed by a max-pooling (stride = 2) to extract the highlighted features (Fig. 1a, Extended Data Fig. 1). Each skip-connection in the residual blocks contains a convolution with a kernel size of 1 to refine the input before adding it to the output. A decoder consists of a transposed convolution, concatenation, and a residual convolution block to reconstruct the output image from the extracted features. To bypass the low-frequency information, we modified the architecture of residual U-Net by replacing the skip-connections between encoder and decoder paths with residual channel attention blocks (CAB; see below). We used three down-samplings and three up-samplings in the encoder and decoder paths, respectively. The initial number of convolutional filters is 64, which is doubled after each pooling in the encoder path while it is halved after each up-sampling in the decoder path. The output layer is a 1×1 convolution.

The second subnetwork is a residual channel attention network (RCAN)^19^, known to be a very deep convolutional neural network for super-resolution image reconstruction. Our RCAN network consists of 3 residual groups (RG) containing 8 CABs and a short skip-connection, a convolution layer, and a long skip connection. Each CAB consists of a convolution block with 64 channels, a global average pooling, a channel down-scaling convolution layer (filter size = 4), followed by a LeakyReLU and a channel upscaling convolution layer. Its output is passed through a sigmoid activation function and is used to rescale the input through multiplication. The upscaling module in the original RCAN was removed since the input and output in our network have the same shape. The number of residual groups and the filter size can increase to improve the performance at the cost of longer training time. All the convolution kernels have a size of 3 unless specified otherwise.

The input of the RCAN is the output of U-Net concatenated with the original noisy input image. While the RCAN network enhances the resolution of the denoised output by U-Net, the original noisy input guides to prevent the loss of spatial information during training. For 3D UNet-RCAN, all the 2D kernels used for convolutions, poolings, and up-samplings were replaced with three-dimensional vectors.

### Preparation of training dataset

We obtained ∼20 pairs of noisy and high SNR STED images (2,048×2,048 pixels) for each target, from which training-set patches were created. The size of our training set for 2D or 3D networks was >1,200 patches (256×256 pixels) or >900 patches (256×256×16 pixels), respectively. Image normalization was performed on the image stacks such that each patch was normalized to its maximum. To exclude patches containing less information from the training dataset, we calculated the L2 norm of each patch, normalized it to the maximum of the norms of the training dataset, and discarded patches with their normalized norm being smaller than a threshold (0.2∼0.4).

### Registration of noisy and ground-truth images

An essential step before training a denoising model is a *xy*-drift correction between noisy and high SNR STED images. This was realized by calculating the cross-correlation of each pair of noisy and high SNR images in the Fourier domain. The drift between images was obtained by the maximum of the cross-correlation. We implemented this algorithm in MATLAB and applied it to the dataset before training our network.

### Preparation of semi-synthetic training dataset

Since Poisson noise is dominant in fast STED imaging, a semi-synthetic dataset can be generated by adding noise to high SNR STED data to make it resemble noisy STED data. We first adjusted the intensity of a high SNR STED image by multiplying it with a coefficient λ. We generated a random Poisson number at each pixel by using the pixel value as a random variable such that synthetic noisy images were prepared. We compared a histogram of this image with that of a noisy STED image obtained by fast STED imaging with a certain pixel dwell time and found the value of λ, which minimized the mean squared error (MSE) between the histograms. We used the average value of λ by repeating this procedure 5 times. It is important to discard the first bin of histograms and normalize them to their maximum before calculating MSE. We used this approach for denoising fast 3D STED images (Fig. X) and live-cell mitochondrial dynamics (Fig. X).

### Training UNet-RCAN

We optimized a loss function which is a weighted summation of Charbonnier loss (*L*_*char*_) and edge loss (*L*_*edge*_)^23^. The Charbonnier loss and edge loss are defined as:

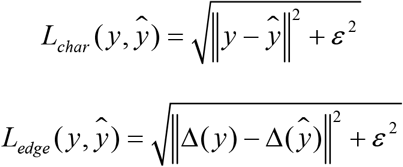

where y is the ground-truth image, ý is the predicted image, Δ is the Laplacian operator, and *ε* is a constant set to 10^−3^ (ref 19). The Laplacian operator was implemented as a convolution of an image with a Laplacian filter. Combining the two loss functions prevents the smoothing effect that usually happens when training with the MSE loss function and ensures the reconstruction of super-resolution images (ref). The total loss function for training UNet-RCAN is defined as:

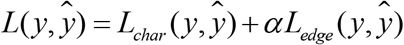

where α is the weight parameter which is empirically set to 0.05 (ref 19).

We implemented our model using Keras^24^ with a Tensorflow backend^25^ in Python. We used an Adam optimizer with the default parameters to minimize our loss function. The initial learning rate was set to 1×10^−4,^ which is scheduled to change using the cosine annealing method^26^. We chose this method to prevent the model from converging to a local minimum. The batch size for training was set to 1 to prevent our GPU memory (12GB) from filling. The models were trained for 200 epochs (2D) and 100 epochs (3D) on an NVIDIA GeForce RTX 3080 Ti graphics card. The training time for 2D and 3D models were approximately 8 h and 24 h, respectively (Supplementary Table 1).

For STED power dependence experiments, each UNet-RCAN network was trained for denoising images of β-tubulin labeled with STAR635P, which were captured at 0, 10, 20, 40, 50, and 70% of STED power. To verify that our prediction results follow the scaling law of STED microscopy^27^, we compared the resolution of our predicted results with that of 20 nm crimson beads (ThermoFisher) at different STED powers.

### Training other networks

We implemented CARE in Keras, according to https://github.com/CSBDeep/CSBDeep. The model was trained on 1,200 patches with 256×256 pixels, a batch size of 16, and an initial learning rate of 4×10^−4^. 2D-RCAN^19^ was implemented using Keras with 5 residual groups (RG) and 10 channel attention blocks (CAB) within each RG. The RG filter shape was set to 64, and the CAB filter shape was set to 4. The model was trained on 1,200 patches with 256×256 pixels, a batch size of 1, and an initial learning rate of 1×10^−4^. We trained these models by optimizing the MSE loss function using an Adam optimizer.

To compare our two-step prediction approach to one-step prediction by modified U-Net or RCAN as described earlier, each network was separately trained for denoising STED images of microtubules. The U-Net filter shape was chosen to be [32,64,128], and the RCAN filter shape was set to 32 with 3 residual groups and 8 channel attention blocks. The filter shape of channel attention blocks was set to 4.

### Quantitative assessment of prediction results

To evaluate the predicted results, a test set of 10 different images with a shape of 2,048×2,048 was analyzed by peak signal-to-noise ratio (PSNR) and multi-scale structural similarity index (MS-SSIM) using the built-in functions of TensorFlow. Spatial resolution was quantified by an ImageJ plug-in for decorrelation analysis^28^ (Radius min = 0, Radius max = 1, Nr = 50, Ng = 10). The average and standard deviation of these parameters for all the predictions results are calculated and displayed in Extended Data Fig. 5 and Supplementary Tables 2-4.

### STED microscopes

Confocal and STED images were acquired using a Leica SP8 3X STED with an oil objective (HC PL APO 100x/NA1.4, Leica) or an Abberior STED Expert Line with an oil objective (UPLXAPO 100x/NA1.45, Olympus). The depletion beams were pulsed lasers emitting at 775 nm. For the Leica system, the excitation power was set to 20%, and the images were detected with HyD detectors (a gain value of 20). We used a resonant scanning mode with a line speed of 8 kHz. The gating window was set to 0.4-12 ns. For 3D STED imaging, the z-STED was activated with 50% of the STED power. 3 line-averaging (*Δt* = 0.054 μs) or 128 line-averaging (*Δt* = 2.3 μs) was applied for collecting the noisy or the ground-truth data. For high-throughput imaging, an *xy* grid of 31×24 STED images with 20% overlap between tiles was obtained with 3-line-averaging and the Leica autofocusing system. For the Abberior system, the excitation power was set to 4.5%, and the images were detected with avalanche photodiodes. The gating window was set to 0.75 – 8 ns. We used a quad galvo scanner with a pixel time of 1 μs. Live-cell STED imaging was performed at room temperature. For details on the imaging conditions, please see Supplementary Table 5.

### Denoising live-cell STED imaging on mitochondrial dynamics

We generated a semi-synthetic dataset as described above (Supplementary Fig. 1 and 2). Briefly, we used high SNR STED images (*Δt* = 90 μs) as ground-truth and generated noisy inputs which have comparable SNR to fast live-cell STED images of mitochondria (*Δt* = 1 μs). The trained network was applied to the noisy live-cell videos to restore high SNR STED time-lapse images.

### Photobleaching assessment

To compare the photobleaching effects of conventional STED and fast STED imaging with deep learning, five different field-of-views were imaged for each imaging modality. Denoising was performed by UNet-RCAN on the fast STED data. To obtain the photobleaching curves, the L2 norm of the noisy data was calculated over the frames and normalized to the maximum of norms. This vector was applied to the prediction results normalized to their maximum over the frames. The average intensity of each frame for denoised fast STED and conventional STED images was plotted as a function of frame number (Fig.2b,2d).

### Cell culture

For imaging immunolabeled samples, U2OS cells (human bone osteosarcoma, HTB-96, ATCC) were grown in McCoy’s 5A medium (ATCC) supplemented with 10% fetal bovine serum (FBS, Sigma-Aldrich, F2442) and 1% penicillin-streptomycin (ThermoFisher), and seeded on coverslips 2-3 days before experiments. For imaging mitochondria dynamics, HeLa cells^29^ were grown in Dulbecco’s Modified Eagle Medium (DMEM) with glutaMAX and 4.5 g/L glucose (ThermoFisher), 1% (v/v) penicillin-streptomycin (Sigma-Aldrich), 1 mM sodium pyruvate (Sigma-Aldrich), and 10% (v/v) FBS (Merck Millipore) at 37°C in a 5% CO_2_ incubator. The cells were seeded in glass-bottom dishes (ibidi GmbH) one day prior to imaging.

### Immunofluorescence labeling

U2OS cells were fixed with 4% paraformaldehyde (Electron Microscopy Sciences, 15710) and 0.2% glutaraldehyde (Electron Microscopy Sciences, 16019) in phosphate buffered saline (PBS) for 15 min at room temperature, then washed in PBS. After incubation in 0.1% (w/v) sodium borohydride (Sigma-Aldrich) for 10 minutes, the cells were washed with PBS three times, followed by blocking with 3% bovine serum albumin (BSA, ThermoFisher) in PBS and permeabilization with 0.5% Triton-X 100 (Sigma-Aldrich) in PBS. When labeling microtubules, the cells were fixed with 0.6% paraformaldehyde, 0.1% glutaraldehyde, and 0.25% Triton-X 100 in PBS for 1 min at 37°C. The cells were incubated in a primary antibody solution diluted to a final concentration of 2.5 μg/mL in PBS overnight at 4°C. After washing three times in PBS, the cells were incubated in a secondary antibody solution diluted to a final concentration of 5 μg/mL in PBS overnight at 4°C. After washing three times in PBS, a cover slip was mounted on a glass microscope slide using Mowiol (Sigma-Aldrich)^30^. Immunolabeling reagents are listed in Supplementary Table 6.

### Mitochondria labeling in living cells

The HeLa cells were stained with DMEM containing 250 nM PK Mito Orange (Confocal.nl)^31^ for 40 min, followed by three washing steps in DMEM. The cells were kept in the incubator for 1hr to remove unbound dyes. The culture medium was replaced with HEPES buffered DMEM containing 4.5 g/L glucose, L-glutamine, and 25 mM HEPES (ThermoFisher). The cells were then imaged at room temperature using the Abberior system.

### Preparation of membrane system for imaging vesicle dynamics

Giant unilamellar vesicles made of 1-palmitoyl-2-oleoyl-glycero-3-phosphocholine (POPC) and cholesterol (2:1 molar ratio) were prepared following the electroformation method^8^. A lipid mixture (5 μL, 1 g/L) dissolved in chloroform were spread onto platinum wires mounted in a custom made polytetrafluoroethylene chamber. The lipid mixture was dried with a gentle stream of N_2_ and subsequently submerged in a 300 mM sucrose buffer. The wires were connected to a function generator. A 10 Hz 2.0 V sine wave was applied for 1h, with the frequency being reduced to 2 Hz for an extra 30 minutes. Supported lipid bilayers made of 1,2-dioleoyl-sn-glycero-3-phosphocholine (DOPC), 1,2-dioleoyl-snglycero-3-phosphoethanolamine (DOPE), and 1,2-dioleoyl-sn-glycero-3phospho-L-serine (DOPS) (molar ratio 4:3:3) were prepared following the spin coating method.

The lipid mixture (25 μL of 1 g/L) dissolved in chloroform:methanol (1:1 volume ratio) were spin-coated (30 s, 3000 rpm) on plasma treated coverslips (#1.5). The coverslips were then mounted on AttoFluor chambers (ThermoFisher), hydrated in HEPES-buffered saline, and cleaned 10 times. The giant vesicles were then transferred to the supported lipid bilayer chamber and after labelling with 200 nM of the exchangeable membrane dye NR4A^8^ and let 15 minutes to settle. To promote membrane fusion 10 mM CaCl_2_ dissolved in HEPES-buffered saline were added.

### Membrane dynamics imaging

Images were acquired on an Abberior Expert Line system^8^ equipped with a UPlanSApo 60×/1.2 water immersion objective lens. Depletion in the *z* direction strongly depended on the correct adjustment of the objective lens correction collar. NR4A was excited with a 561 nm laser with a 10 μW laser power at the sample plane. Depletion was achieved using a 775 nm (40 MHz) with a power of 300 mW at the sample plane.

## Data availability

Data may be obtained from the authors upon reasonable request.

## Code availability

The codes, sample data and instruction guide are available at the GitHub repository (https://github.com/vebrahimi1990/UNet_RCAN_Denoising.git).

## Acknowledgments

We thank A.S. Belmont for valuable discussions, and C. Ullal, J. Cha and N. Urban for generously permitting the use of their STED systems. We thank C. Ullal and A. Husain for critically reading our manuscript. This work was based in part on data recorded using an instrument whose acquisition was supported by the US National Science Foundation (1725984). This work was supported by the US National Institutes of Health (R35GM138039 and U01DK127422 to K.Y.H.), the European Research Council Advanced Grant (ERC AdG No. 835102) and the DFG-funded CRC 1286 (project A05). P. Carravilla received funding from the European Commission Horizon 2020 Marie Skłodowska Curie programme (H2020-MSCA-IF-2019-ST project 892232 FILM-HIV).

## Author contributions

V.E. and K.Y.H. conceived the project. V.E., J.K. and K.Y.H. designed the experiments. V.E., J.K., T.S. and P.C. performed the experiments. C.E., S.J. and K.Y.H. supervised research. V.E. and K.Y.H. wrote the paper with input from all authors.

## Competing financial interests

The authors declare no competing financial interests.

**Extended Data Fig. 1.**
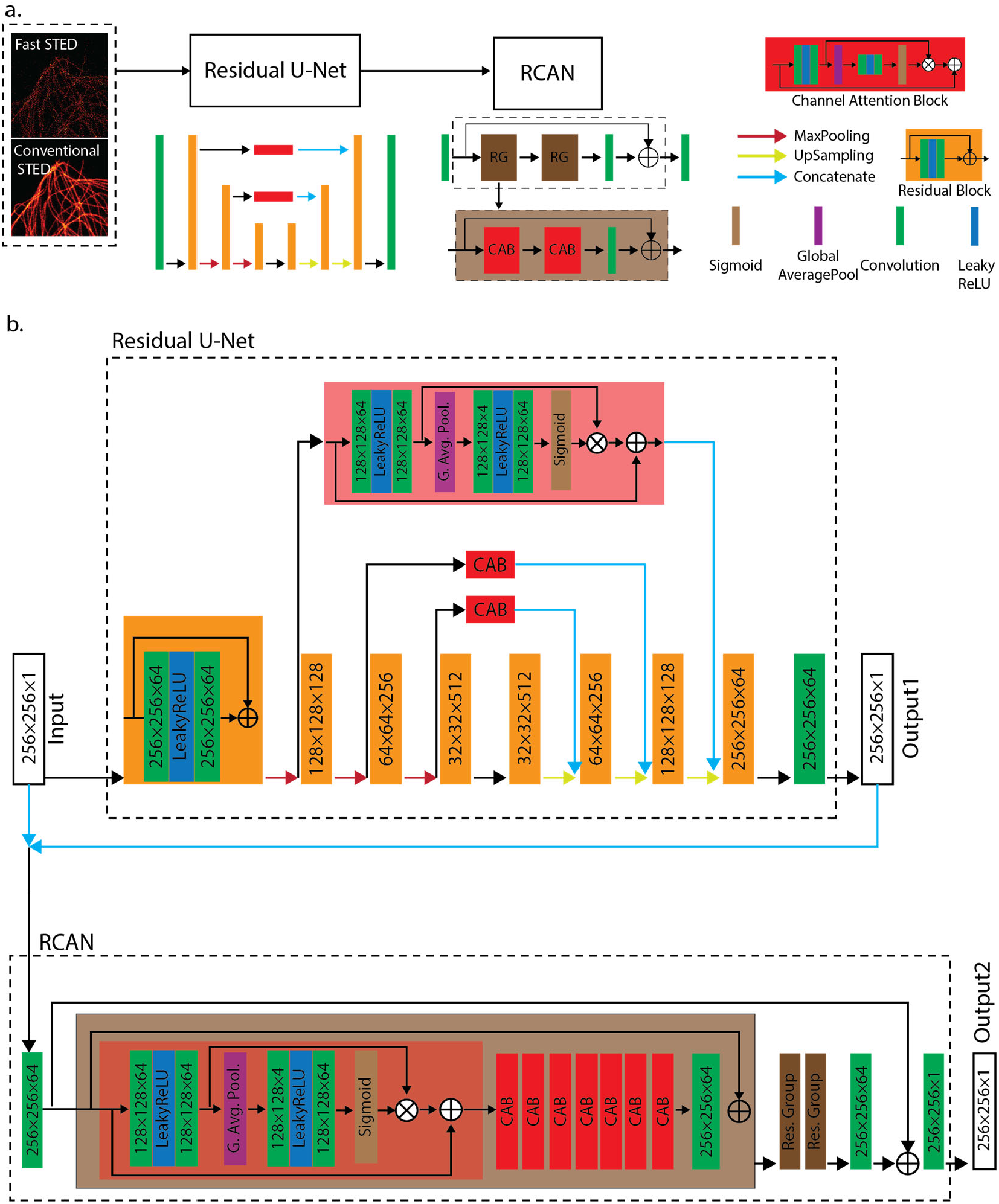
The architecture of UNet-RCAN for denoising fast STED imaging data. (a) The schematic of two-step prediction with UNet-RCAN. (b) Detailed architecture of UNet-RCAN. CAB, channel attention block; G. Avg. Pool., global average pooling; Res. Group, residual group. The model has one input and two outputs. Both output1 and output2 contribute to the loss function.

**Extended Data Fig. 2.**
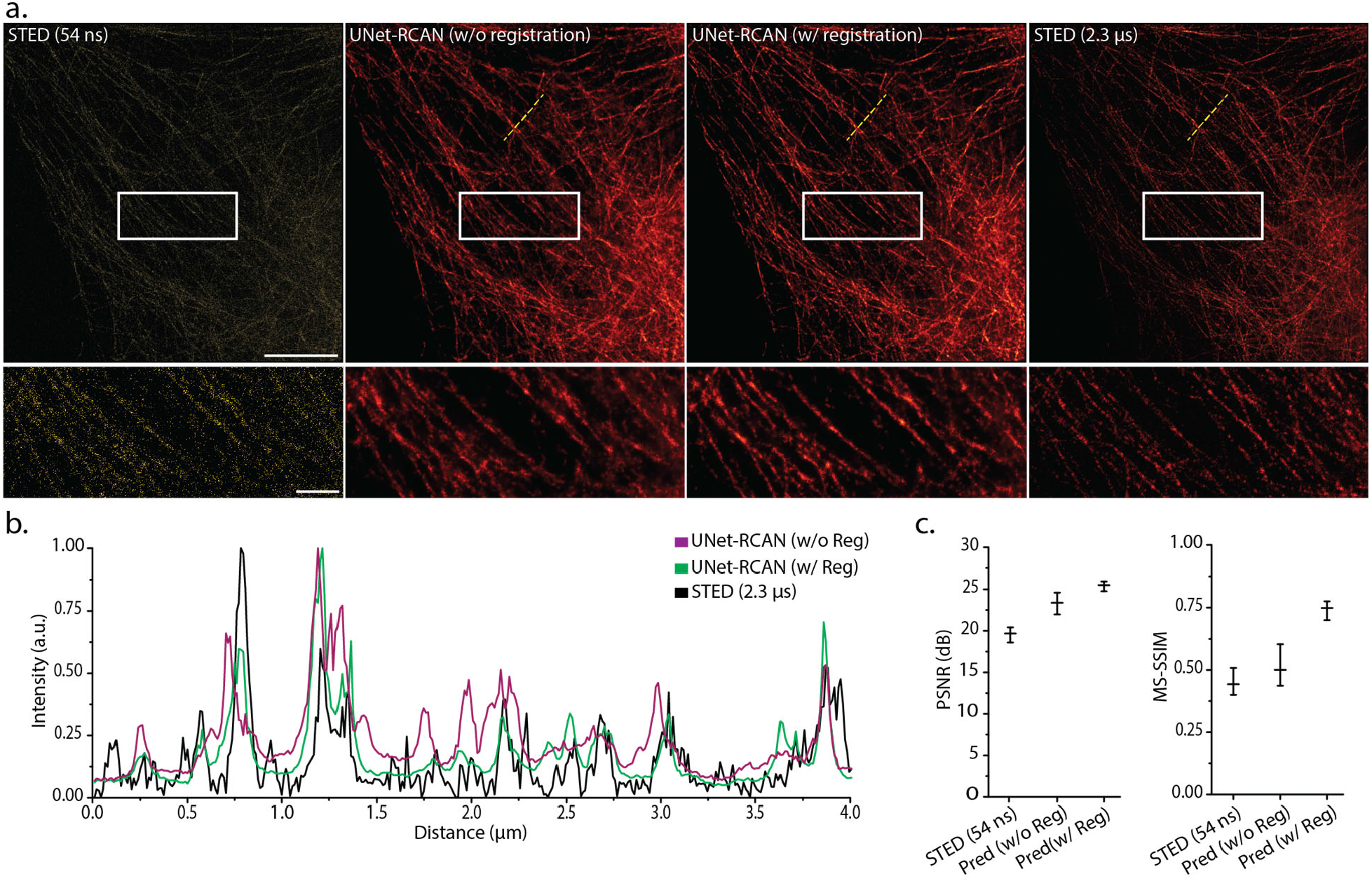
Registration of noisy input and ground-truth STED images. Registration improves prediction accuracy. Predicted STED images (a), line profiles (b), and quantitative assessment (c) by UNet-RCAN with and without registration of the noisy and ground-truth images by drift correction. Line profiles were measured along the yellow dashed lines. Scale bars, 5 μm and 1 μm (magnified regions).

**Extended Data Fig. 3.**
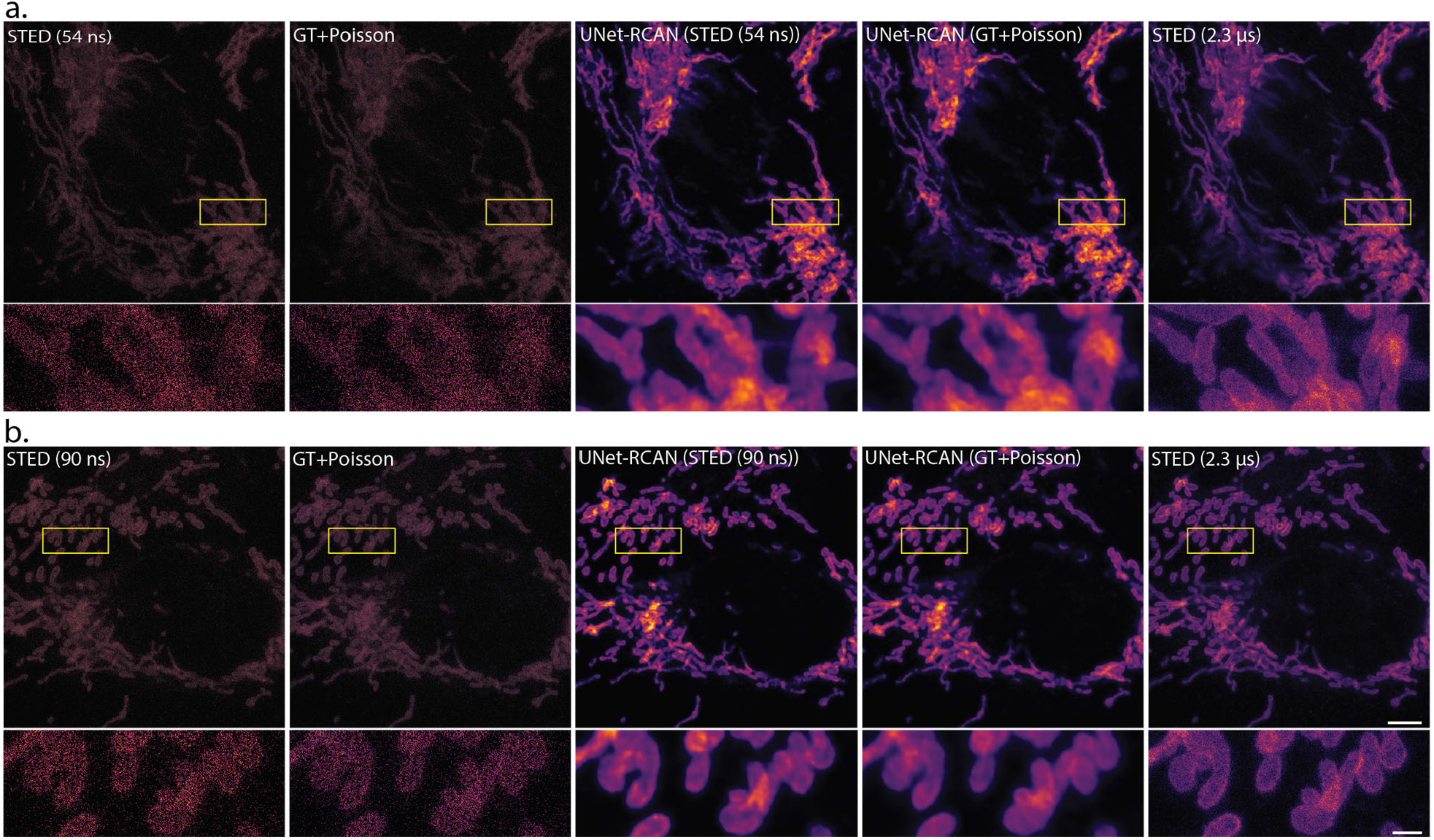
Comparison of semi-synthetic data and sequentially acquired noisy data. The predicted STED results from semi-synthetic data show similar results to those of well registered pairs of images. Different levels of SNR were tested, i.e., with a pixel time of 54 ns (a) or 90 ns (b) by adding Poisson noise to STED data with a pixel time of 2.3 μs. STED images of TOM20 labeled with Atto647N in U2OS cells. Scale bars, 5 μm and 1 μm (magnified regions).

**Extended Data Fig. 4.**
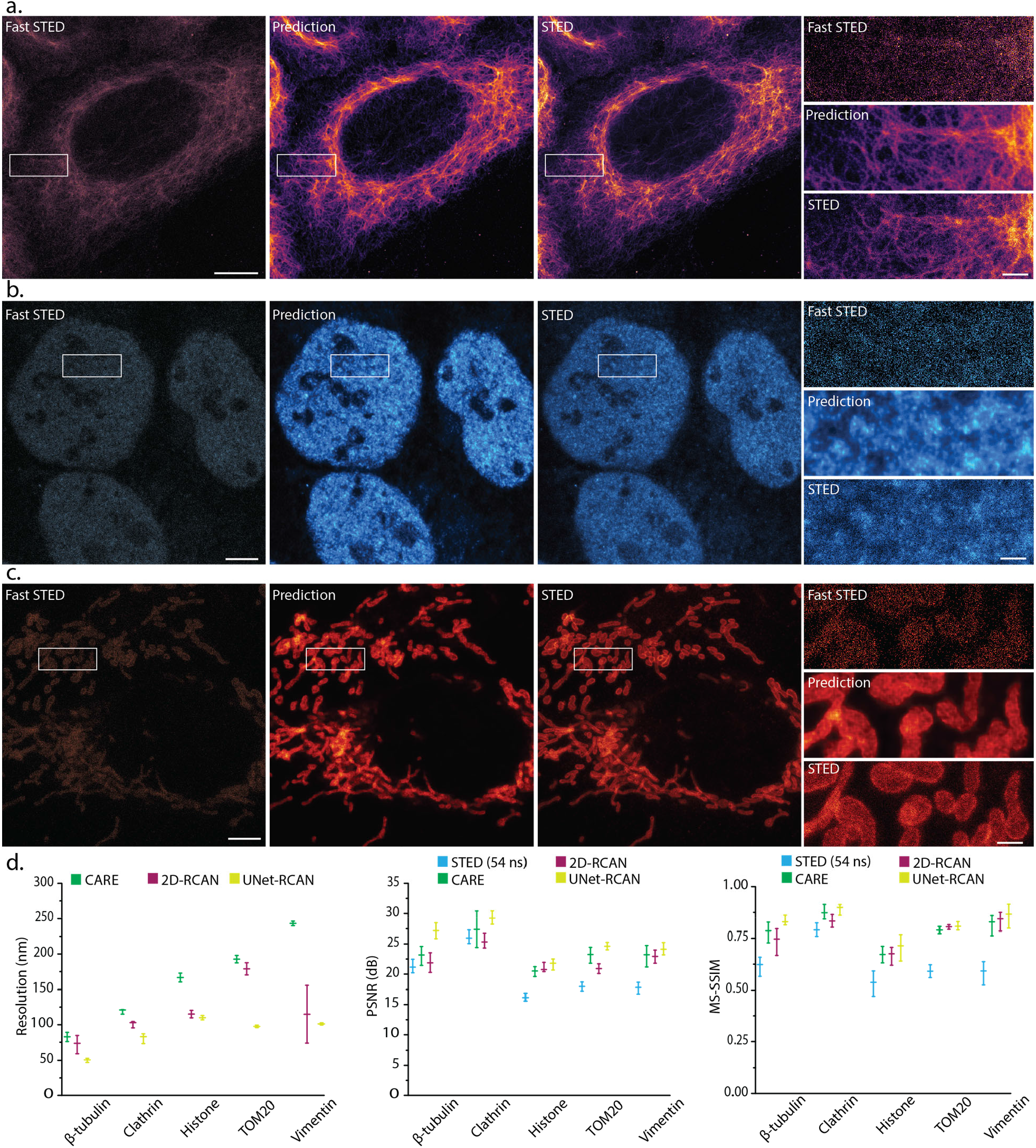
Denoising results of 2D-STED images by UNet-RCAN on subcellular structures and comparison of the performances of UNet-RCAN, CARE, and 2D-RCAN. (a) Vimentin labeled with STAR635P, (b) Histone (H3K9ac) labeled with Atto647N, and (c) TOM20 labeled with Atto647N in fixed U2OS cells. Scale bars, 5 μm and 1 μm (magnified regions). (d) Five different markers were used for testing: β-tubulin (STAR635P), clathrin (STAR580), histone (Atto647N), TOM20 (Atto647N), and vimentin (STAR635P). The resolution was obtained by decorrelation analysis. Mean and standard deviation are displayed (*n* = 10).

**Extended Data Fig. 5.**
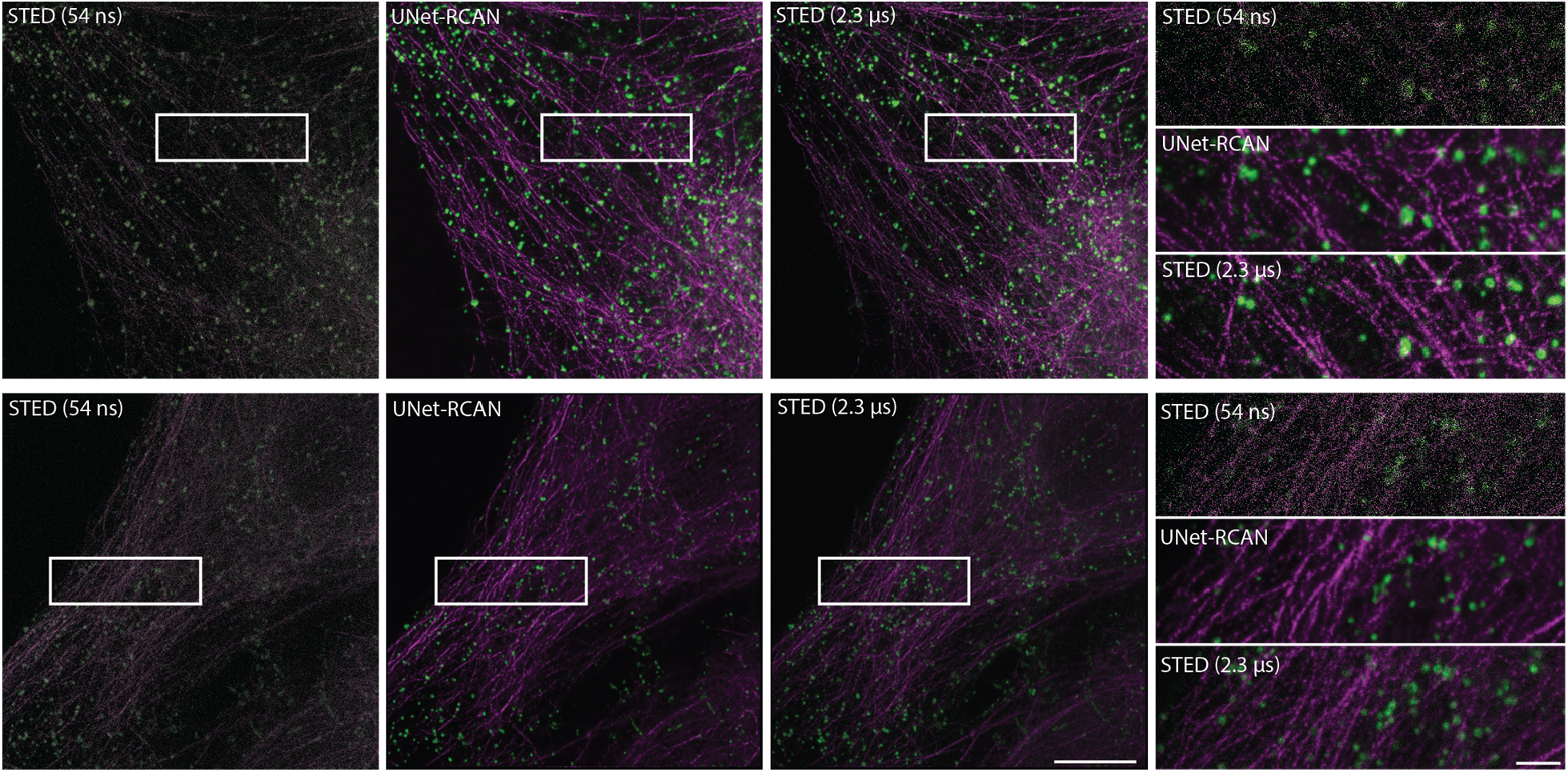
Denoising two-color fast STED imaging with UNet-RCAN. STED images of β-tubulin (STAR635P, magenta) and clathrin (STAR580, green) in U2OS cells with a pixel time of 54 ns. The ground-truth STED images were captured with a pixel time of 2.3 μs. Scale bar, 5 μm and 1 μm (magnified regions).

**Extended Data Fig. 6.**
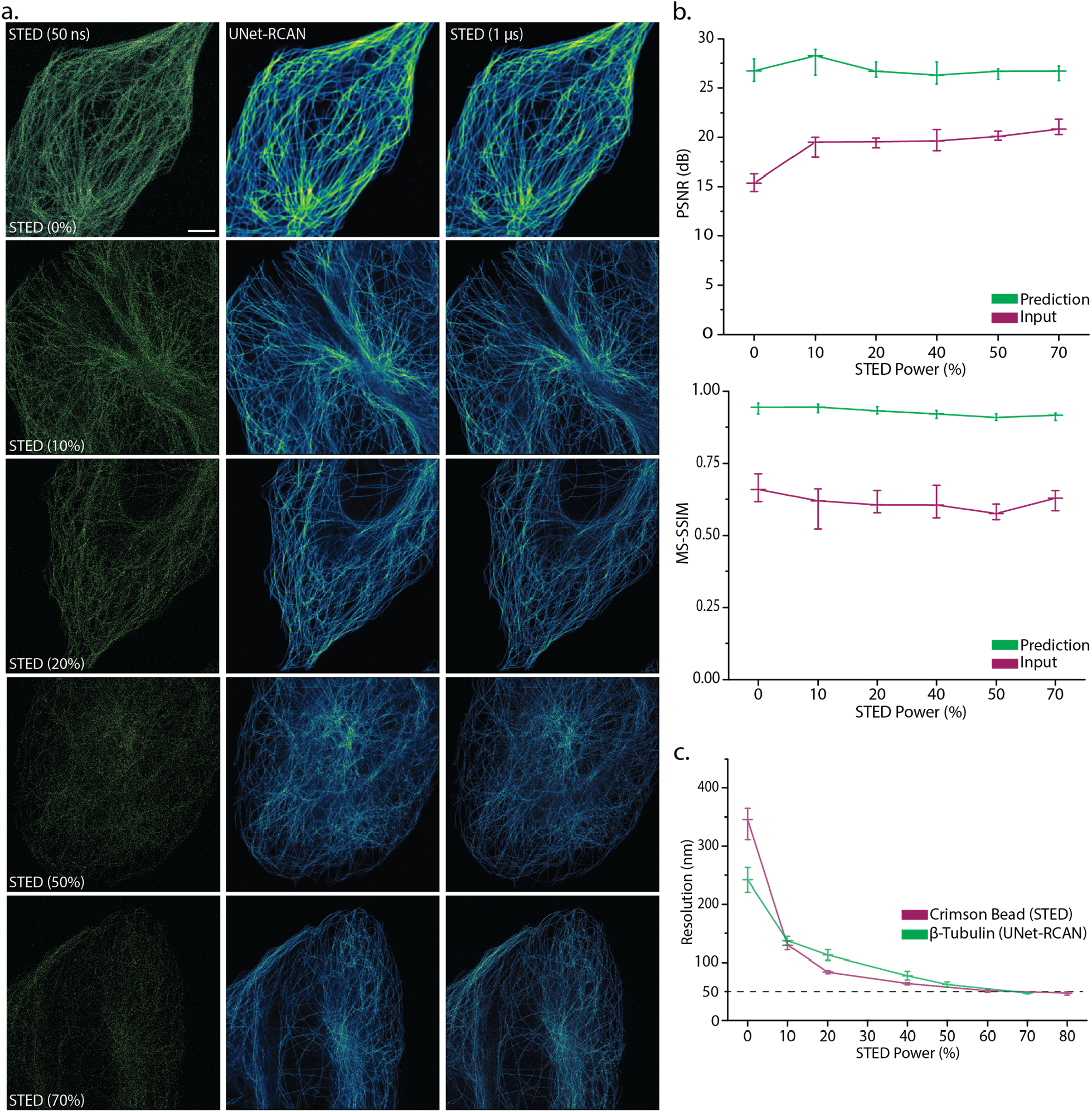
STED power dependence. (a) Noisy (*Δt* = 50 ns), ground-truth (*Δt* = 1 μs) and UNet-RCAN STED images on β-tubulin (STAR635P) in U2OS cells. Six different datasets were generated by STED imaging with 0, 10, 20, 40, 50, and 70% of the STED power. (b) MS-SSIM and PSNR analysis for denoising results at each STED power. Mean and standard deviation are displayed (*n* = 8). (c) The resolution of the prediction results (green) corresponds well with that of 20 nm crimson beads (magenta). The resolution was calculated by decorrelation analysis. Mean and standard error of mean are displayed (*n* = 8). Scale bar, 5 μm.

**Extended Data Fig. 7.**
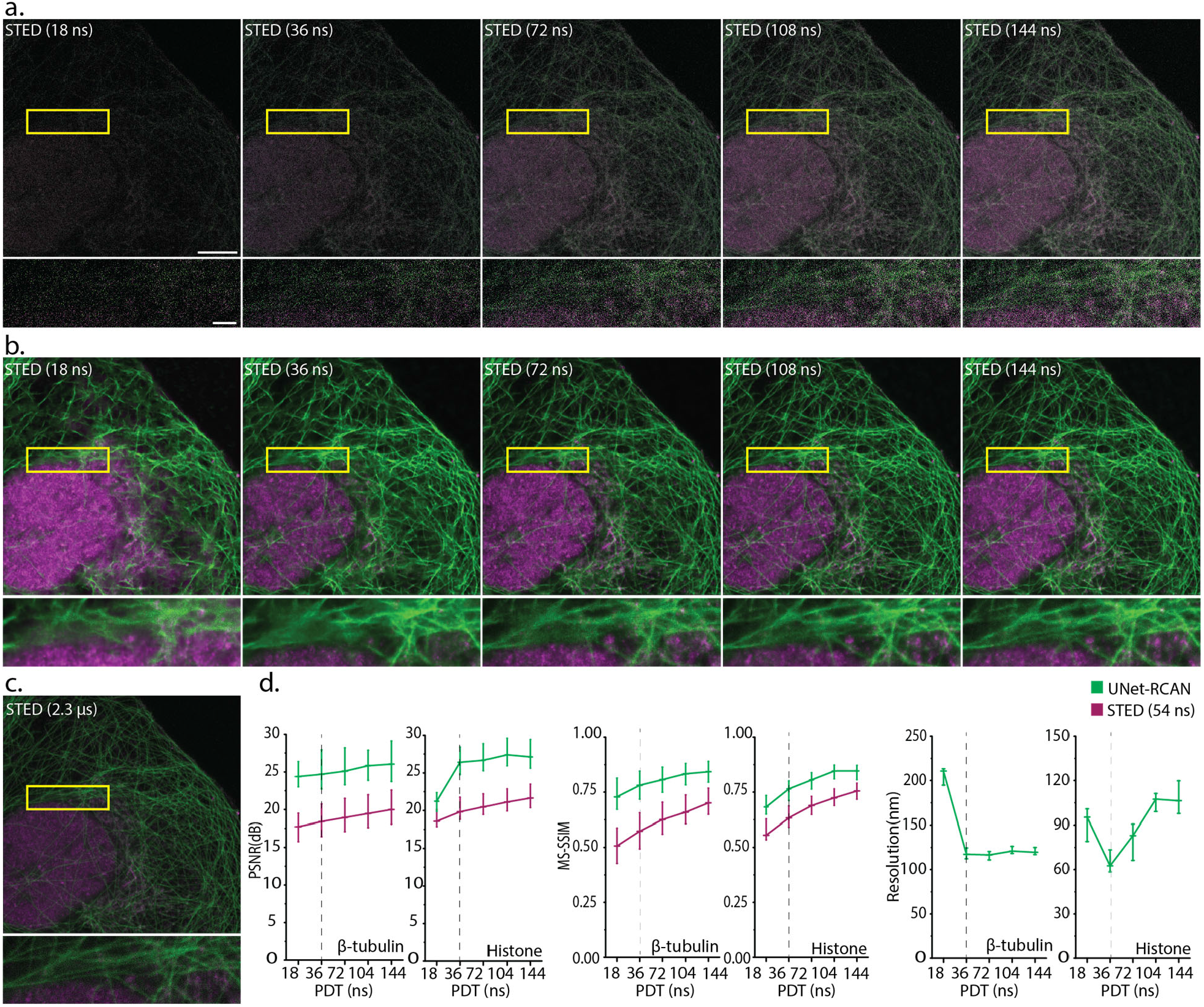
UNet-RCAN can denoise STED images with different SNR levels. (a) Noisy two-color STED images of β-tubulin (STAR580, green) and histone (Atto647N, magenta) with pixel times of 18, 36, 72, 104, and 144 ns. (b) Prediction results of (a) by UNet-RCAN. (c) Ground-truth STED image with a pixel time of 2.3 μs. (d) PSNR, MS-SSIM, and resolution analysis for each pixel dwell time. Mean and standard deviation are displayed (*n* = 10). Scale bars, 5 μm and 1 μm (magnified regions).

**Extended Data Fig. 8.**
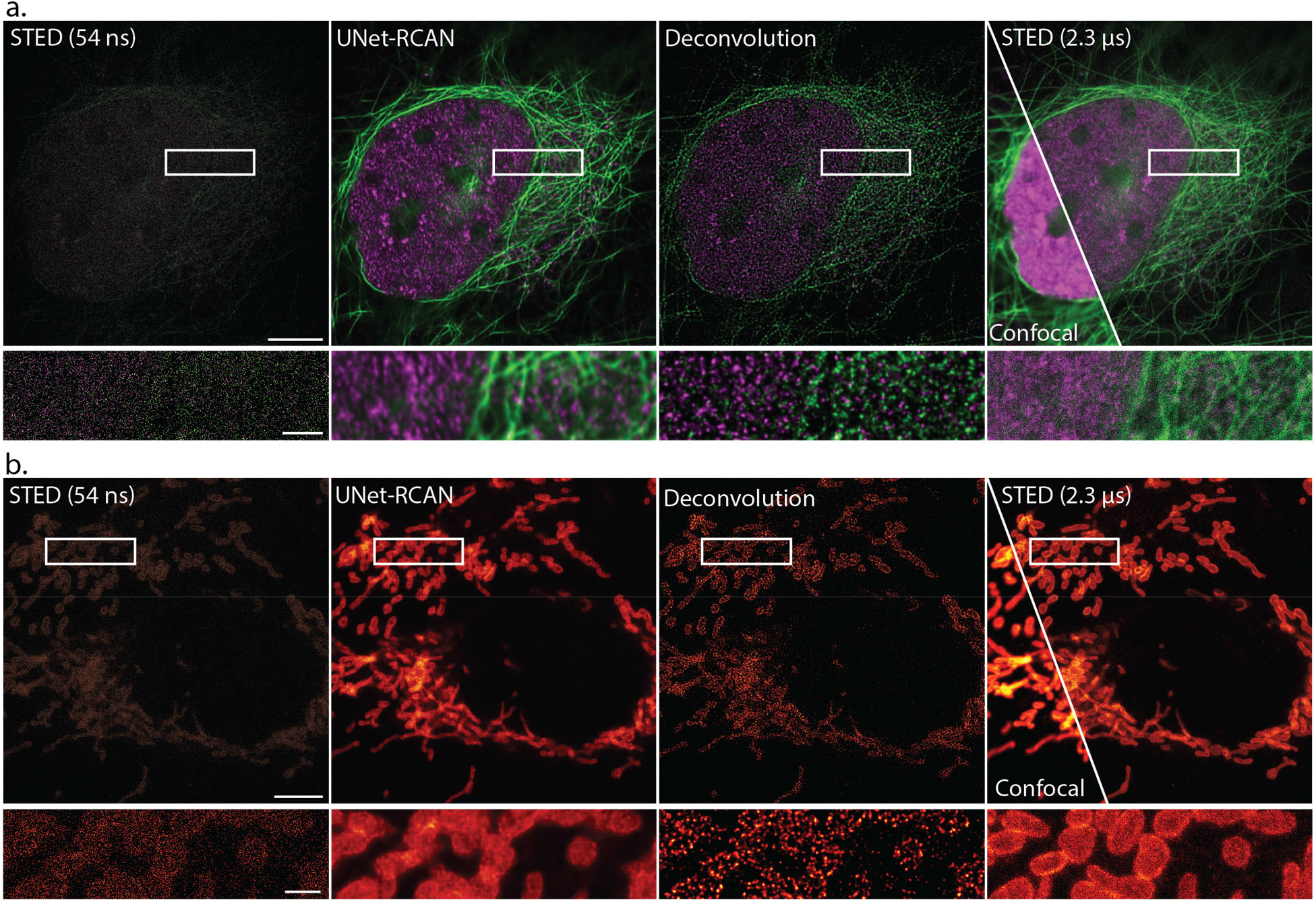
Comparison of denoising by UNet-RCAN and deconvolution. Denoising fast STED imaging of (a) β-tubulin (STAR580, green) and histone (Atto647N, magenta), and (b) TOM20 (Atto647N) in U2UOS cells. Separate denoising tasks were performed by UNet-RCAN and deconvolution on noisy STED data captured with a pixel time of 54 ns. Deconvolution was performed with Huygens software (See Methods). Scale bars, 5 μm and 1 μm (magnified regions).

**Extended Data Fig. 9.**
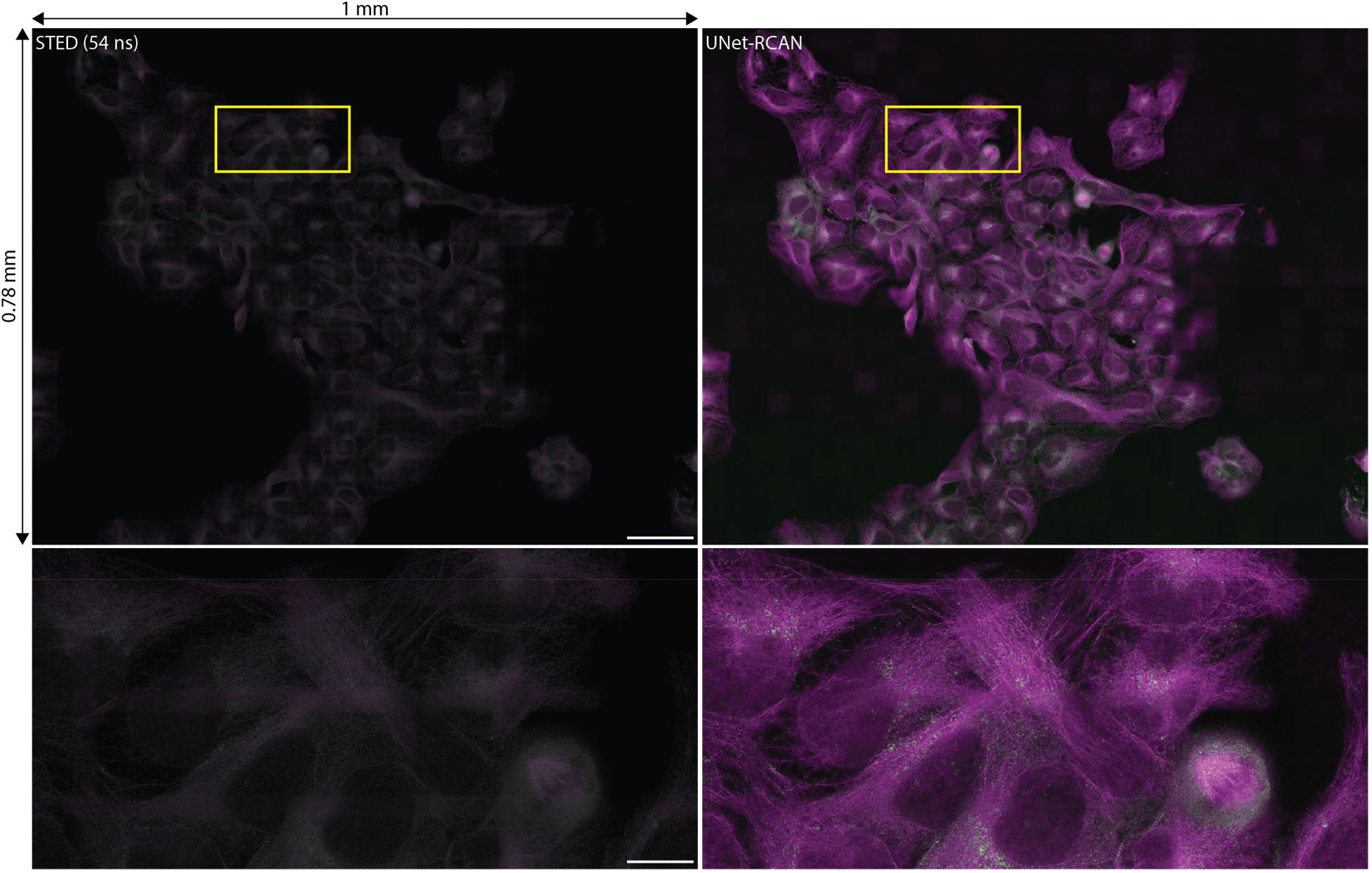
High-throughput STED imaging with UNet-RCAN. STED images of β-tubulin (STAR635P, magenta) and clathrin (STAR580, green) in U2OS cells with a pixel time of 54 ns. A grid of 31×24 images was denoised with UNet-RCAN and stitched together. Scale bars, 100 μm and 20 μm (magnified regions).

**Extended Data Fig. 10.**
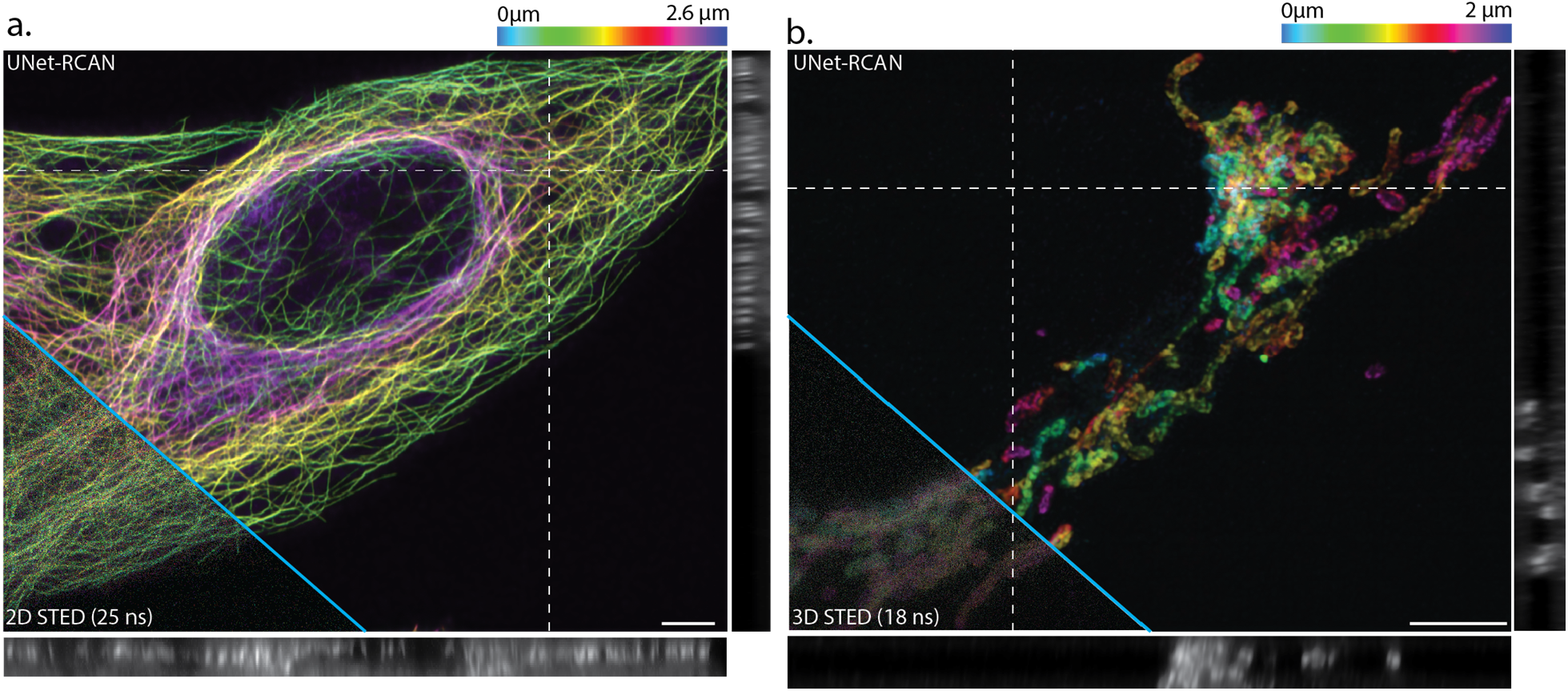
Prediction by 3D-UNet-RCAN on 2D and 3D STED imaging. 3D-UNet-RCAN prediction results for a noisy z-stack of (**a**) 2D-STED imaging of β-tubulin (STAR635P) and (**b**) 3D-STED imaging of TOM20 (Atto647N) in U2OS cells. The z pixel sizes are 170 nm and 65 nm, respectively. Scale bars, 5 μm.

