## Supplementary Information for "Deep learning enables fast, gentle STED microscopy"

**Supplementary Videos**

**Supplementary video 1.** Time-lapse STED imaging of mitochondria dynamics with a pixel time of 90 µs. HeLa cells were labeled with PK Mito Orange.

**Supplementary video 2.** Fast deep-learning STED imaging of mitochondria dynamics with a pixel time of 1 µs. HeLa cells were labeled with PK Mito Orange.

**Supplementary video 3.** Two-color live-cell deep-learning STED imaging of mitochondria (green) and ER (magenta) in HeLa cells with a pixel time of 1 µs. Mitochondria was labeled with PK Mito Orange, and ER was labeled with SiR-Halo.

**Supplementary video 4.** Deep-learning live-cell STED imaging with deconvolution. COS-7 cells were labeled with PK Mito Orange.

**Supplementary video 5.** Denoising fast 3D STED *xz* imaging of giant unilamellar vesicles (GUV) labeled with NR4A with a pixel time of 2 µs.

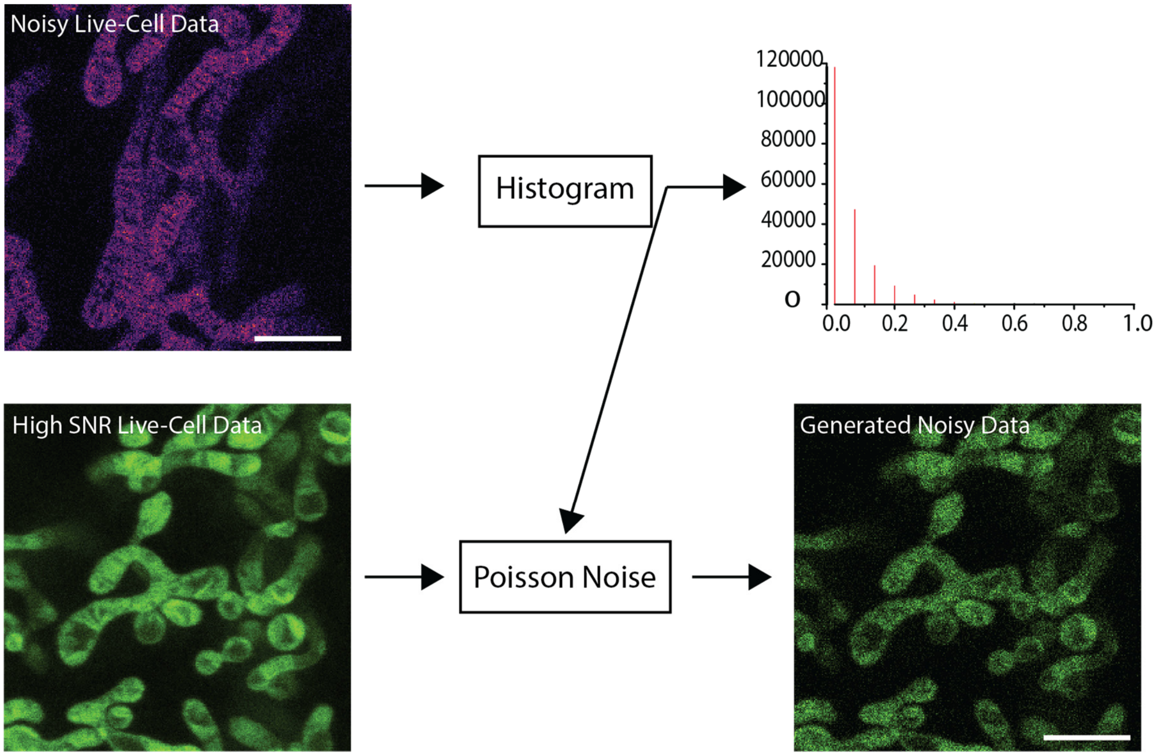


**Supplementary Fig. 1, Semi-synthetic dataset generation.** Poisson noise was applied to the high SNR STED images of cristae labeled with PK Mito Orange in HeLa cells to generate a pair of noisy and high SNR data for training UNet-RCAN. The amount of Poisson noise was adjusted such that the intensity histogram of the generated noisy data resembles that of the noisy live cell STED data (See Methods). Scale bars, 2µm.

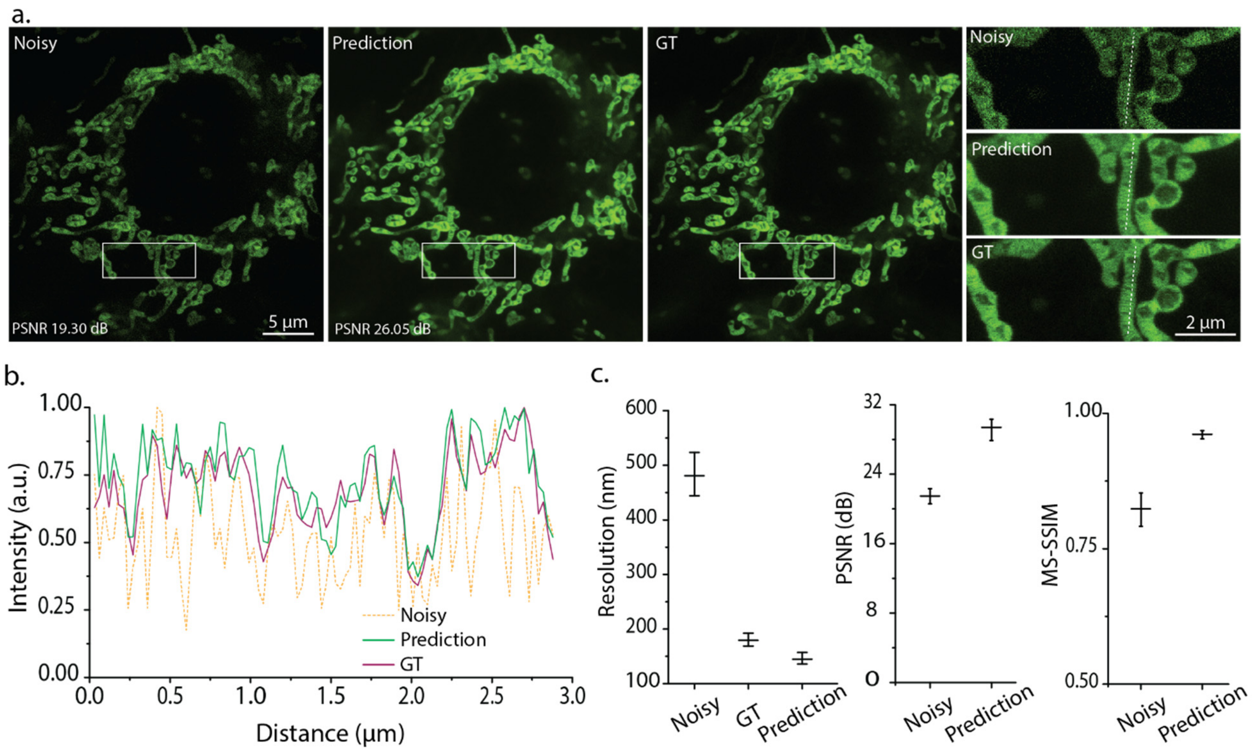


**Supplementary Fig. 2, Denoising performance of UNet-RCAN on the semi-synthetic dataset.** (a) Denoising results of cristae labeled with PK Mito Orange in HeLa cells. The GT data was captured with a dwelling time of 90 µs. The noisy data was generated by adding Poisson noise. The prediction is the denoising result by UNet-RCAN. (b) Line profiles of noisy, prediction, and GT data along the dashed lines in (a). (c) Resolution analysis by decorrelation, PSNR, and MS-SSIM calculations were performed on the prediction results by UNet-RCAN. Mean and standard deviation are displayed (*n* = 10).

### Supplementary Note 1, Comparisons with other deep learning approaches

In cross-modality image restoration, a diffraction-limited confocal image is transformed into a super-resolved STED image with resolution enhancement by a deep convolutional neural network[1]. Different network architectures could perform this image transformation, such as generative adversarial networks (GAN)[2] or residual channel attention networks (RCAN)[3]. A transformation between confocal and STED imaging modalities at least requires a 3~5-fold resolution enhancement, i.e., from 250 nm to 50 nm; however, the cross-modality deep learning approaches have proven to be limited by a factor of 2-2.5 in terms of resolution enhancement[3]. Moreover, a lack of enough information often leads to exhibit artifacts.

Unlike the cross-modality image transformation, denoising is performed on noisy but super-resolved STED images to improve SNR. In denoising, the input data contains more information in terms of spatial resolution. This can help to reduce artifact generation and improve resolution enhancement. In Figs.1f-h, we showed that denoising STED data clearly outperforms the cross-modality approach perceptually and according to the image quality assessment parameters.

Two popular network architectures suitable for denoising super-resolution data are UNet and RCAN. A UNet learns the features in an image dataset through convolutional layers and multiple down-sampling and up-sampling layers. Although the UNet effectively denoises diffraction-limited imaging data such as widefield images, its output does not reliably preserve high-frequency information. This is likely due to the fact that there is no mechanism to prioritize high-frequency information. Moreover, in UNet, a final image is reconstructed through downsampling and upsampling layers rather than applying the convolutional filters on the original noisy super-resolved data.

On the other hand, RCAN contains channel attention blocks and several skip connections, which help prioritize and maintain high-frequency information in super-resolution image reconstruction. Moreover, in RCAN, final super-resolved images are restored by applying filters on the original noisy data. This lowers the possibility of missing high-frequency information. However, our STED denoising results with RCAN show that although its final result is superior to UNet in terms of resolution, it generates more high-frequency artifacts, which may be due to its CAB building blocks, especially when the input SNR is extremely poor.

We showed that by combining UNet and RCAN, denoising could be effectively performed on fast STED data while we can maintain the super-resolution and prevent high-frequency artifact generation.

### Supplementary Tables

**Supplementary Table 1.** Parameters and training time for CARE, 2D-RCAN, and UNet-RCAN.

| | CARE | 2D-RCAN | UNet-RCAN |
|---|---|---|---|
| # Iterations per epoch | 70 | 1,080 | 1,080 |
| Batch size | 16 | 1 | 1 |
| Patch size | 256×256 | 256×256 | 256×256 |
| Epochs | 200 | 200 | 200 |
| Number of parameters | 3,790,850 | 3,944,073 | 16,684,270 |
| Training time | 1 h 17 m | 11 h 40 m | 8 h 10 m |

**Supplementary Table 2.** Performance comparison chart of UNet-RCAN, CARE, and 2D-RCAN in terms of SNR.

| | Noisy | CARE | 2D-RCAN | UNet-RCAN |
|---|---|---|---|---|
| β-tubulin | 21.3±1.1 dB | 23.0±1.6 dB | 22.0±1.6 dB | 27.2±1.3 dB |
| Clathrin | 26.2±1.1 dB | 27.4±3.0 dB | 25.6±1.2 dB | 29.3±1.1 dB |
| Histone | 16.2±0.6 dB | 20.4±0.8 dB | 21.2±0.8 dB | 21.6±0.9 dB |
| TOM20 | 18.0±0.7 dB | 23.1±1.3 dB | 20.9±0.8 dB | 24.6±0.6 dB |
| Vimentin | 17.7±0.9 dB | 23.0±1.8 dB | 23.0±1.1 dB | 24.2±1.1 dB |

**Supplementary Table 3.** Performance comparison chart of UNet-RCAN, CARE, and 2D-RCAN in terms of similarity.

| | Noisy | CARE | 2D-RCAN | UNet-RCAN |
|---|---|---|---|---|
| β-tubulin | 0.61±0.04 | 0.77±0.05 | 0.73±0.06 | 0.83±0.02 |
| Clathrin | 0.79±0.03 | 0.87±0.03 | 0.83±0.03 | 0.88±0.02 |
| Histone | 0.53±0.06 | 0.67±0.04 | 0.66±0.04 | 0.70±0.06 |
| TOM20 | 0.59±0.03 | 0.79±0.02 | 0.80±0.01 | 0.81±0.02 |
| Vimentin | 0.58±0.05 | 0.81±0.05 | 0.83±0.04 | 0.85±0.06 |

**Supplementary Table 4.** Performance comparison chart of UNet-RCAN, CARE, and 2D-RCAN in terms of resolution measured by decorrelation analysis.

| | CARE | 2D-RCAN | UNet-RCAN |
|---|---|---|---|
| β-tubulin | 80±3 nm | 71±3 nm | 51±1 nm |
| Clathrin | 122±5 nm | 103±6 nm | 81±4 nm |
| Histone | 166±2 nm | 115±2 nm | 110±1 nm |
| TOM20 | 193±2 nm | 179±3 nm | 98±2 nm |
| Vimentin | 243±1 nm | 115±13 nm | 101±1 nm |

**Supplementary Table 5.** Acquisition settings of STED imaging.

| Figures | | |
|---|---|---|
| 1b, 2a<br>ED 2a,4a,5, 9 | Exc. power = 20%<br>STED power = 50%<br>Resonant scanning<br>Gating: 0.4-12 ns<br>Leica STED | Fluorophore: STAR635P, $\lambda_{exc}$ = 635 nm, $\lambda_{STED}$ = 775 nm<br>Pixel time: 0.054 μs (noisy) and 2.3 μs (ground-truth) |
| 2a | | Fluorophore: Alexa 594, $\lambda_{exc}$ = 594 nm, $\lambda_{STED}$ = 775 nm<br>Pixel time: 0.054 μs (noisy) and 2.3 μs (ground-truth) |
| ED 10a | | Fluorophore: STAR635P, $\lambda_{exc}$ = 635 nm, $\lambda_{STED}$ = 775 nm<br>Pixel time: 0.025 μs (noisy) and 1μs (ground-truth) |
| 1f<br>ED 3a, 4b, 4c, 8a, 8b | | Fluorophore: Atto647N, $\lambda_{exc}$ = 647 nm, $\lambda_{STED}$ = 775 nm<br>Pixel time: 0.054 μs (noisy) and 2.3 μs (ground-truth) |
| ED 3b | | Fluorophore: Atto647N, $\lambda_{exc}$ = 647 nm, $\lambda_{STED}$ = 775 nm<br>Pixel time: 0.090 μs (noisy) and 2.3 μs (ground-truth) |
| ED 5, 8a, 9 | | Fluorophore: STAR580, $\lambda_{exc}$ = 580 nm, $\lambda_{STED}$ = 775 nm<br>Pixel time: 0.054 μs (noisy) and 2.3 μs (ground-truth) |
| ED 7a, 7c | | Fluorophore: Atto647N, $\lambda_{exc}$ = 647 nm, $\lambda_{STED}$ = 775 nm<br>Pixel time: [0.018,0.036,0.072,0.108,0.144] μs (noisy) and 2.3 μs (ground-truth) |
| ED 7a, 7c | | Fluorophore: STAR580, $\lambda_{exc}$ = 580 nm, $\lambda_{STED}$ = 775 nm<br>Pixel time: [0.018,0.036,0.072,0.108,0.144] μs (noisy) and 2.3 μs (ground-truth) |
| ED 6a | Exc. power = 20%<br>STED power = [0%,10%,20%,50%,70%],<br>Resonant scanning<br>Gating: 0.4-12 ns<br>Leica STED | Fluorophore: STAR635P, $\lambda_{exc}$ = 635 nm, $\lambda_{STED}$ = 775 nm<br>Pixel time: 0.050 μs (noisy) and 1.0 μs (ground-truth) |
| 2g, ED 10b | Exc. power = 20%<br>2D-STED power = 50%<br>z-STED power = 50%<br>Resonant scanning<br>Gating: 0.4-12 ns | Fluorophore: Atto647N, $\lambda_{exc}$ = 635 nm. $\lambda_{STED}$ = 775 nm<br>Pixel time: 0.018 μs (noisy) and 2.3 μs (ground-truth) |

| | | |
|---|---|---|
| | Leica STED | |
| 2c, 2d<br>SI 2a | Exc. power = 4.5%<br>STED power = 22%<br>Galvo scanning<br>Gating: 0.75-8 ns<br>Abberior STED | Fluorophore: PK Mito Orange, $\lambda_{exc}$ = 561 nm, $\lambda_{STED}$ = 775 nm<br>Pixel time: 1 μs (noisy) |
| 2h | Exc. power = 35%<br>2D-STED power = 0%<br>z-STED power = 100%<br>Galvo scanning<br>Gating: 0 ns<br>Abberior STED | Fluorophore: NR4A , $\lambda_{exc}$ = 561 nm, $\lambda_{STED}$ = 775 nm<br>Pixel time: 2 μs (noisy) and 20 μs (ground-truth) |

**Supplementary Table 6.** Immunolabeling conditions.

| Figures | Primary antibody | Secondary antibody | Fluorophore |
|---|---|---|---|
| 1b, 2a<br>ED 2a, 5, 6a, 9 | Monoclonal Anti-β-Tubulin antibody produced in mouse, Sigma-Aldrich, T5293 | Fab Fragment Goat Anti-Mouse IgG1, Jackson ImmunoResearch, 115-007-185 | Abberior STAR 635P |
| ED 7a, 7c, 8a | Monoclonal Anti-β-Tubulin antibody produced in mouse, Sigma-Aldrich, T5293 | Fab Fragment Goat Anti-Mouse IgG1, Jackson ImmunoResearch, 115-007-185 | Abberior STAR 580 |
| ED 5, 9 | Anti-Clathrin heavy chain antibody (ab21679) | Fab Fragment Goat Anti-Rabbit IgG, Jackson ImmunoResearch, 111-007-008 | Abberior STAR 580 |
| 1f<br>ED 4b, 7a, 7c, 8a,<br>SI 4a | Anti-acetyl-Histone H3 (Lys9) in rabbit, Sigma-Aldrich, 07-352 | Rabbit IgG (H&L) Antibody ATTO 647N Conjugated Pre-Adsorbed, ROCKLAND, 611-156-122 | Atto 647N |
| 2a | Anti-acetyl-Histone H3 (Lys9) in rabbit, Sigma-Aldrich, 07-352 | Alexa Fluor® 594 AffiniPure $F(ab')_2$ Fragment Goat Anti-Rabbit IgG (H+L) | Alexa Fluor 594 |
| 2g<br>ED 3a, 3b, 4c, 10b | Anti-TOMM20 antibody - Mitochondrial Marker, abcam, ab78547 | Rabbit IgG (H&L) Antibody ATTO 647N Conjugated Pre-Adsorbed, ROCKLAND, 611-156-122 | Atto 647N |

| | | | |
|---|---|---|---|
| ED 4a | Anti-Vimentin antibody, Mouse monoclonal (V6389-200UL) | Fab Fragment Goat Anti-Mouse IgG1, Jackson ImmunoResearch, 115-007-185 | Abberior STAR 635P |
